# Simple phenotypic tests to improve accuracy in screening chromosomal and plasmid-mediated colistin resistance in gram-negative bacilli

**DOI:** 10.1101/2020.07.06.190850

**Authors:** Fernando Pasteran, Diego Danze, Alejandra Menocal, Carla Cabrera, Ignacio Castillo, Ezequiel Albornoz, Celeste Lucero, Melina Rapoport, Paola Ceriana, Alejandra Corso

## Abstract

CLSI and EUCAST recommends that only broth microdilution (BMD) should be used for routine colistin susceptibility testing, however, it could be difficult to perform in resource-poor settings. The purpose of this study was to evaluate the accuracy of an agar spot test (COL-AS) and a colistin drop test (COL-DT) as compared to BMD. COL-AS and COL-DT were challenged with a collection of 271 Gram-negative bacilli clinical isolates: 195 Enterobacterales (including 63 mcr-1 positive strains), 37 Acinetobacter spp. and 39 P. aeruginosa. For COL-AS, 3.0µg/ml (final concentration) of colistin was added to a Mueller-Hinton agar plate and subsequently swabbed with a 0.5 McFarland suspension of tested strain within 1cm2 spot. For COL-DT, 10µl of a 16µg/ml colistin solution was dripped on the surface of a Mueller-Hinton agar plate, previously inoculated with a lawn of tested strain (0.5 McFarland). Colistin solution was made either, by dissolving powder or by disk elution in CA-MHB. Overall, 141/271 (52%) isolates were categorized as colistin resistant by reference BMD. COL-AS yielded a categorical agreement (CA) of 95.5% compared to BMD, with 0.7% very major errors and 3.8% major errors. COL-DR yielded a CA of 96.2% compared to BMD, with 0.7% and 0% very major errors and 3.1% and 3.8% major errors, for colistin powder and disk elution solutions, respectively. Most major errors occurred for mcr-1 producing strains with MICs that fluctuated from 2 to 4 μg/ml. In conclusion, we developed and validated methods suited to the systematic screening of resistance to colistin in gram negative bacilli.

**Clinical relevance:** colistin continues to be one of the last-line therapeutic options to treat carbapenemase-producing gram negative bacilli. The BMD reference methodology, the only one accepted by current standards for evaluating colistin sensitivity, is difficult to implement in laboratories from low-resource countries. In this work, we propose two simple methodologies to screen for colistin resistance, with a performance equivalent to the reference method in detecting resistance to colistin, both of plasmid (mcr) and chromosomal nature. Furthermore, the methods validated here allowed a better identification of those producers of mcr producer with borderline MICs. These screening tests can be routinely performed in addition to the tests currently in use. showed long stability during storage and some of them do not require colistin powder as the source of antibiotic, an important limitation in low-resource countries.

## BACKGROUND

Colistin (polymyxin E) and polymyxin B are old polycationic peptides that have regained popularity as last resort treatments to face the worldwide emergence of multidrug-resistant gram-negative bacteria (1). Even though acquired resistance to polymyxins was seen only occasionally in the past, this is becoming more common because of the renewed interest in the polymyxins (2). The recent identification of *mcr* genes, coding for plasmid-mediated resistance mechanisms, suggests that resistance to polymyxins might not only be mediated by chromosomal mutations but also spread by horizontal transmission, which warranted immediate worldwide attention (3, 4). These genes have been found primarily in *Escherichia coli* from human, animal, food and environmental samples (5, 6).

In 2020, CLSI issued colistin breakpoints for *Enterobacterales, Pseudomonas aeruginosa and Acinetobacter baumannii* complex but only considering intermediate (≤ 2 µg/ml) and resistant categories (≥ 4 µg/ml), based on the limited clinical efficacy of this drug (7). While the clinical breakpoints issued by EUCAST for the same group of bacteria included susceptible (≤ 2 µg/ml) and resistant (> 2 µg/ml) categories (8). Both, CLSI and EUCAST guidelines recommend routine colistin susceptibility testing by estimation of MIC, being the broth microdilution-BMD- (without the addition of surfactant) the reference standard method (7, 8). However, there are important limitations for MIC estimation by BMD in several facilities, especially in resource-poor settings, being a labor-intensive method and the accessibility to colistin sulfate the main ones.

Additionally, determination of the MIC by other methods, such as gradient strips or semi-automatic equipment, as Phoenix or Vitek, have been discouraged by EUCAST (http://www.eucast.org/ast_of_bacteria/warnings/#c13111, last accessed 2^nd^ July, 2020) or questioned in their performance due to very major errors in recent publications, respectively (9-13). The disk diffusion method is used in many clinical laboratories either as their primary test method or as a backup method for agents not included on their laboratory’s routine test panels. However, measurements of *in vitro* colistin resistance by disk diffusion is not reliable because large molecular weight antimicrobials, as polymyxins, diffuse slowly into agar, resulting in small differences in the size of inhibition zones between susceptible and non-susceptible isolates (12, 14).

Thus, the dilemma of how to best perform colistin susceptibility testing is a pressing concern for clinical laboratories, even greater now with the threat of the dissemination of carbapenemase producers. The objective of this study was to develop and compare the accuracy of user-friendly colistin testing methods to BMD for screening colistin resistance. Tests developed in this work included the Colistin Agar Spot (COL-AS, addition of a unique concentration of a colistin solution on an agar plate) and the colistin/polymyxin B drop test (COL/POL-DT, application of a single drop of a polymyxin solution on the agar surface). These user-friendly tests resulted suitable for the systematic screening of resistance to colistin in gram-negative bacilli. (Part of this work was presented at the 28^th^ European Congress of Clinical Microbiology & Infectious Diseases; 2018).

## MATERIALS AND METHODS

### Collection and characterization of strains

A panel of previously characterized and de-identified 271 gram-negative clinical isolates, collected from a variety of sources (August 2012 to August 2017), belonging to the repository collection of the NRL, were used in this study. Strains were stored at - 70°C. Panel isolates were previously identified using classical biochemical tests or matrix-assisted laser desorption ionization–time of flight mass spectrometry (MALDI-TOF MS) (16). Species distribution was as follow: 37 *Acinetobacter* spp. (33 A. *baumannii*, 1 *A. pittii*, 1 *A. haemolyticus*, 1 *A. junii*, 1 *A. ursingii*), 39 *Pseudomonas aeruginosa*, 195 *Enterobacterales* (1 *Citrobacter farmeri*, 1 *C. amalonaticus*, 3 *C. freundii*, 16 *Enterobacter cloacae*, 105 *E. coli*, 1 *Klebsiella aerogenes*, 55 *K. pneumoniae*, 5 *K. oxytoca*, 6 *Salmonella* spp.). About 44 (1 *C. freundii*, 5 *Enterobacter spp*, 1 *E. coli*, 3 *K. oxytoca* and 36 *K. pneumoniae*) and 22 (1 *C. freundii*, 3 *E. coli* and 18 *K. pneumoniae*) isolates were KPC or NDM producers, respectively, as determined by PCR (17). Isolates were screened for the presence of *mcr*-1 genes by PCR, as described (18). *mcr*-1 positive *E. coli* isolates were compared by pulsed-field gel electrophoresis (PFGE) of *Xba*I-digested genomic DNA, as described (19).

### Colistin susceptibility tests

Because polymyxins are able to bind to plastic surfaces, for all the tests, antibiotic stock solutions and dilutions were prepared and stored in glass bottles or tubes to limit as much as possible the contact of colistin with plastic. When required, glass pipettes were also utilized. The same batches of cation-adjusted Mueller Hinton broth (CA-MHB), MH agar (MHA) and colistin solutions were used in all the screening test, unless otherwise expressly indicated. All tests were performed on the same working day, for which the same bacterial suspension was used to inoculate the tests. The screening methods were carried-out from fresh (18 to 24 h of incubation) mono-microbial cultures of the strains under study, growth in either plates of blood agar, chocolate agar, CLDE medium, TSA medium or MHA, according to availability.

### 1) Reference colistin susceptibility tests (BMD, macro-dilution and agar dilution)

Duplicate MIC values to colistin were obtained by the recommended BMD method, according to CLSI/EUCAST (8, 20), using in-house untreated 96-well sterile polystyrene microplates (Nunc, Roskilde, Denmark). Colistin sulfate powder (Sigma St. Louis, Missouri, USA) was dissolved in CA-MHB (BD, Franklin Lakes, USA) (stock solution, 128 µg/ml) and log_2_ dilutions were subsequently made to achieve a MIC range of 0.5 to 64 µg/ml. Cation levels (23.7 mg/L calcium and 11.2 mg/L magnesium) were verified by dry phase measurement (VITROS 4600 System, Ortho Clinical Diagnostics) after supplementation. The strains were considered to have acquired resistance to colistin when the MIC was higher than > 2 μg/ml, according to EUCAST standards (8). For colistin-susceptible *mcr* producers, MICs were confirmed by two additional reference dilution methods, a macro-dilution assay and the agar dilution technique (20).

### 2) COL-DT, Colistin Drop Test

The test described by Jouy E. et al. for *E. coli* was modified in the present work to make it suitable for other bacterial species (21). Briefly, we first tested two-fold concentrations of colistin sulfate solutions ranging from 4 to 64 μg/ml (solutions were the same used for BMD) to select the optimal concentration. We used a reduced panel of isolates, which included representative isolates (two susceptible and two colistin resistant, previously determined by BMD) of each *E. coli, Klebsiella pneumoniae, Acinetobacter spp* and *P. aeruginosa* of the panel (Table S1). Next, a single 10 μl drop of each colistin solution was deposited on a MHA plate (Difco) previously swabbed with a 0.5 McFarland inoculum of the strain. Each colistin solution was evaluated in duplicate. The center of each drop was at least 2 cm away from the others. The dripped plates were left for 15 minutes at room temperature (the drop must be completely absorbed before moving the plate), then inverted and incubated for 16 to 18 h at 35 °C. After incubation, the presence or absence of an inhibitory zone was noted after carefully examination with transmitted light. For the purpose of standardization halos were recorded, but the isolate was categorized as colistin susceptible if any zone of inhibition was observed, or colistin resistant when there was no halo. Colonies within the inhibition zone were indicative of resistant subpopulations and the strain was recorded as colistin resistant. Once the conditions of the tests with better performance have been determined (a 16 μg/ml colistin solution made in CA-MHB showed the best performance in the preliminary tests), the entire panel was evaluated as described above.

Subsequently, we tested additional two polymyxin solutions obtained by elution of commercial disks, namely: a 16 μg/ml colistin solution which was prepared by elution of eight 10 μg colistin disks (BD) in 5 ml volume of CA-MHB (BD). The same procedure was performed with eight 300 UI polymyxin B disks (Oxoid) to obtain a 480 UI/ml (equivalent to 0.06 mg/ml) polymyxin solution. Tubes containing disks were incubated at room temperature for at least 30 minutes but not longer than 60 minutes to allow colistin/ polymyxin to elute from the disks, after which disks were aseptically removed. Solutions were sterilized by 0.22 μm filtration (Millipore) before use and kept un-fractioned at 4°C. All solutions were dripped in the same agar plate, with each drop at least 2 cm away from the others. The stability of solutions was tested monthly for 12 months. Repeatability studies (triplicates, on consecutive days) were performed for selected isolates.

### 2) COL-AS, Colistin Agar Spot

In the spot test, the antimicrobial agent was incorporated into the (molten) agar medium (MHA plates) as recommended for the agar dilution method (20), however, unlike agar dilution, the antimicrobial dilution to be tested is limited to the concentration/s that encompass the interpretive breakpoints. For colistin, colistin solutions were added to two batches of molten MHA (BD) to render 2.0 μg/ml and 3.0 μg/ml colistin (final concentration), respectively, in a 20-ml volume plate (100 × 10 mm). These concentrations were chosen because separate the wild-type populations with MIC <= 2 μg/ml from non-wild-type with MIC > 2.0 μg/ml. Plates were kept in sealed bags at 4°C up to 2 months. The excess moisture was removed before use by plate incubation for 5-15 min a 35°C. Subsequently, a sterile cotton swab was dipped in a 0.5 McFarland suspension of the test strain and the excess liquid removed before spot an agar surface area approximately 20 mm in diameter. The same plate allows up to 12 strains to be tested simultaneously. The inoculated plates were left for 15 minutes at room temperature, then inverted and incubated for 16 to 18 h at 35 °C. Plates were examined carefully with transmitted light. A strain was considered colistin resistant if exhibited >1 colony growth and colistin susceptible when there was no-growth at all. The presence of a single colony was considered possible cross-contamination and the test was repeated. The stability of spot plates was tested weekly for 2 months.

Additionally, we evaluated a commercial spot agar (ColTest, Laboratorios Britania, Argentina) that uses another brand of MHA (Britania) and a proprietary concentration of colistin, using the same procedure as the one described for the homemade tests.

### Data analysis

Categorical agreement (CA) represented the rate of isolates grouped in the same susceptibility category by COL-DT or COL-AS compared with BMD, as recommended by the International Organization for Standardization (ISO) standard 20776-2 and the Federal Drug Administration guidance on review criteria for assessment of antimicrobial susceptibility devices (23, 24). Very major errors (VMEs) were isolates that were susceptible to colistin by screening tests and resistant by BMD (MIC > 2 μg/ml); major errors (MEs) were isolates categorized as resistant by screening tests but susceptible by BMD (MIC <= 2 μg/ml). As we did not use intermediate category in our definition, no minor errors were analyzed. VMEs were calculated using the number of resistant isolates as the denominator, and MEs, using the number of susceptible isolates as the denominator.

Agreement and error rates were evaluated by applying the requirements suggested by FDA for susceptibility testing methods (23, 24): a method must exhibit CA of >89.9%, a MEs rate based on the number of susceptible strains tested of < 3%, a VMEs rate of < 2.86%, based on the number of resistant strains tested included in the validation of the respective device (see result section for further explanations), to be considered acceptable.

## RESULTS

### Colistin reference MICs

Overall, 141/271 (52%) isolates were categorized as colistin resistant by reference BMD: 17/37 (46%) *Acinetobacter* spp. (15/33 *A. baumannii*, 1/1 *A. haemolyticus* and 1/1 *A. junii*), 13/39 (33%) *P. aeruginosa* and 111/195 (57%) *Enterobacterales* (1/1 *C. amalonaticus*, 1/1 *E. aerogenes*, 11/16 *E. cloacae*, 69/105 *E. coli*, 20/55 *K. pneumoniae*, 3/5 *K. oxytoca* and 6/6 *Salmonella* spp.) (Table S1). The distributions of colistin MICs determined by BMD are presented in Figure 1. About 63/271 isolates were found to produce *mcr*-1 (1/1 *C. amanolaticus*, 61/106 *E. coli* and 1/57 *K. pneumoniae*). All 61 *E. coli mcr* producers isolates were non-clonal. By BMD, 4/63 (6%) *E. coli mcr* producers repeatedly (triplicates) showed a MIC value below the cut-off value for colistin (MIC 2.0 µg/ml). Interestingly, no other enterobacteria presented a 2.0 µg/ml colistin MIC. The remaining *mcr* producer strains (59/63) showed a MIC > 2 µg/ml, indicating acquired resistance towards colistin. Using a macro-dilution and agar dilution tests, the 4 *mcr*-producing strains categorized as wild-type by BMD, showed repeatedly (triplicates) a MIC one dilution higher, falling within the resistant category. Skipped wells occurred in 5/16 *E. cloacae* isolates (the isolates were categorized as resistant after retesting).

**Figure 1.**
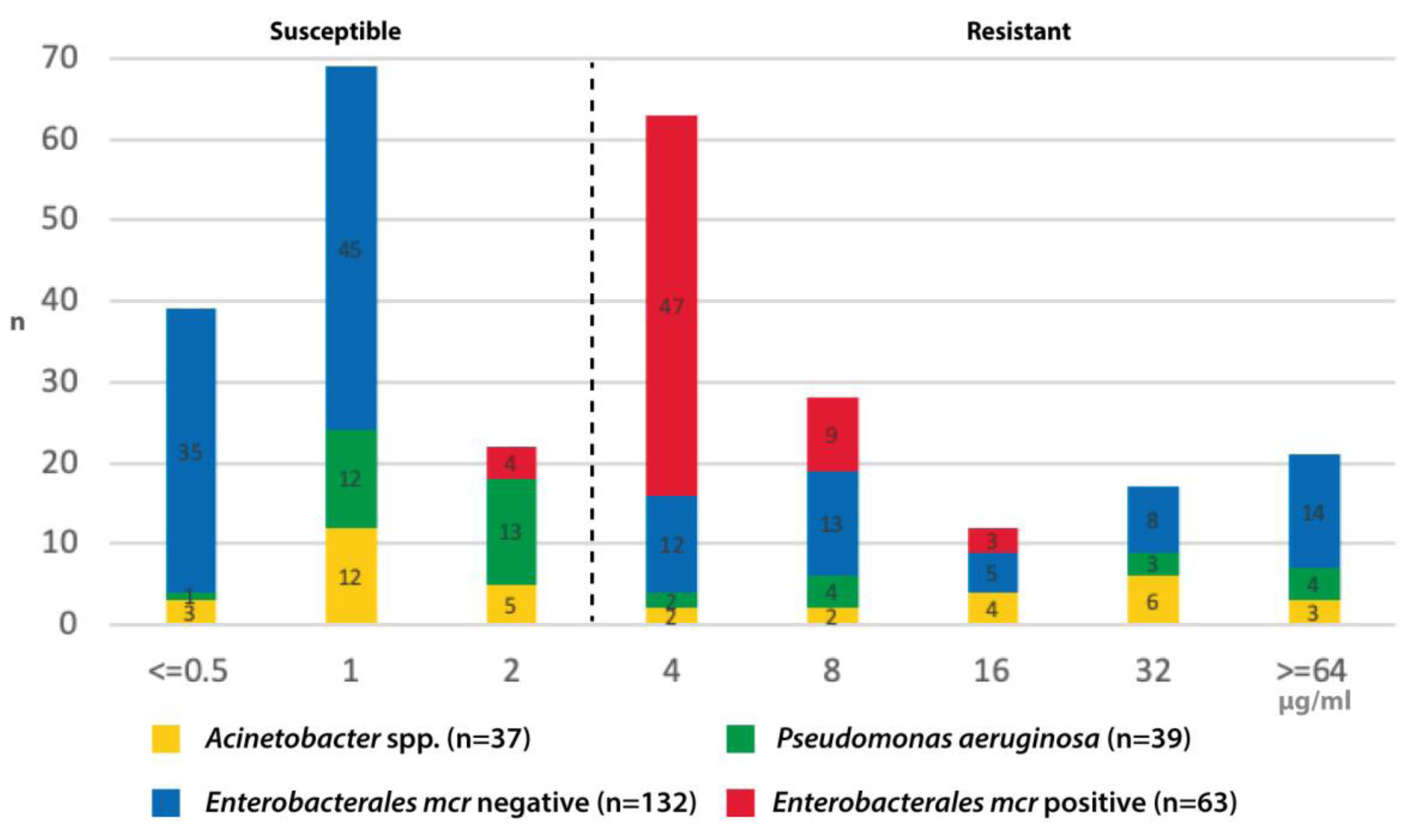
Colistin MIC distributions obtained by BMD. Number of isolates with the indicated colistin MIC value according to bacterial group and the presence of *mcr*-1 gene. The dotted line indicates the breakpoint value that defines susceptible/resistant according to EUCAST standards (Ref. 8).

### Screening tests

#### 1) COL-DT

The first step consisted in determining the optimal concentration of colistin in a 10 μL drop to allow the detection of colistin-resistant bacteria. The minimum colistin concentration in a 10 μL solution to obtain a clear inhibition zone in colistin-susceptible *E. coli* strains was 8 μg/ml, as reported (22). For colistin-susceptible species other than *E. coli*, we found that the minimum colistin concentration in a 10 μL drop to obtain a clear inhibition zone was 16 μg/ml (solution was made using colistin powder dissolved in CA-MHB) (Table 1S).

Using a 16 μg/ml CA-MHB solution, 140/141 of the strains with colistin MIC > 2.0 μg/ml met the criteria for a colistin-resistant drop test result: 130/141 colistin-resistant bacteria grew without an inhibition zone around the colistin drop and 10/141 (4 *Salmonella* spp., 3 *E. cloacae*, 1 *P. aeruginosa* and 2 *E. coli mcr* producer) showed repeatedly well-defined colonies invading the drop zone (Fig. 2; Table 1S). Only one colistin-resistant *P. aeruginosa* isolate was inhibited by the 10 μL drop (zone of 9 mm). The colistin concentration of 16 μg/ml in a 10 μL drop inhibited the growth of 126/130 colistin-susceptible strains (drop zones range/median: 8-11/9.6 mm for *Acinetobacter* spp., 6-12/8.5 mm for *P. aeruginosa* and 8-11/9.6 mm for *Enterobacterales*). Interesting, 4 *E. coli mcr*-1 producers with BMD MICs of 2.0 μg/ml but macro-dilution/agar dilution MICs of 4.0 μg/ml grew without inhibition zone around the drop and were classified as resistant by COL-DT.

**Figure 2.**
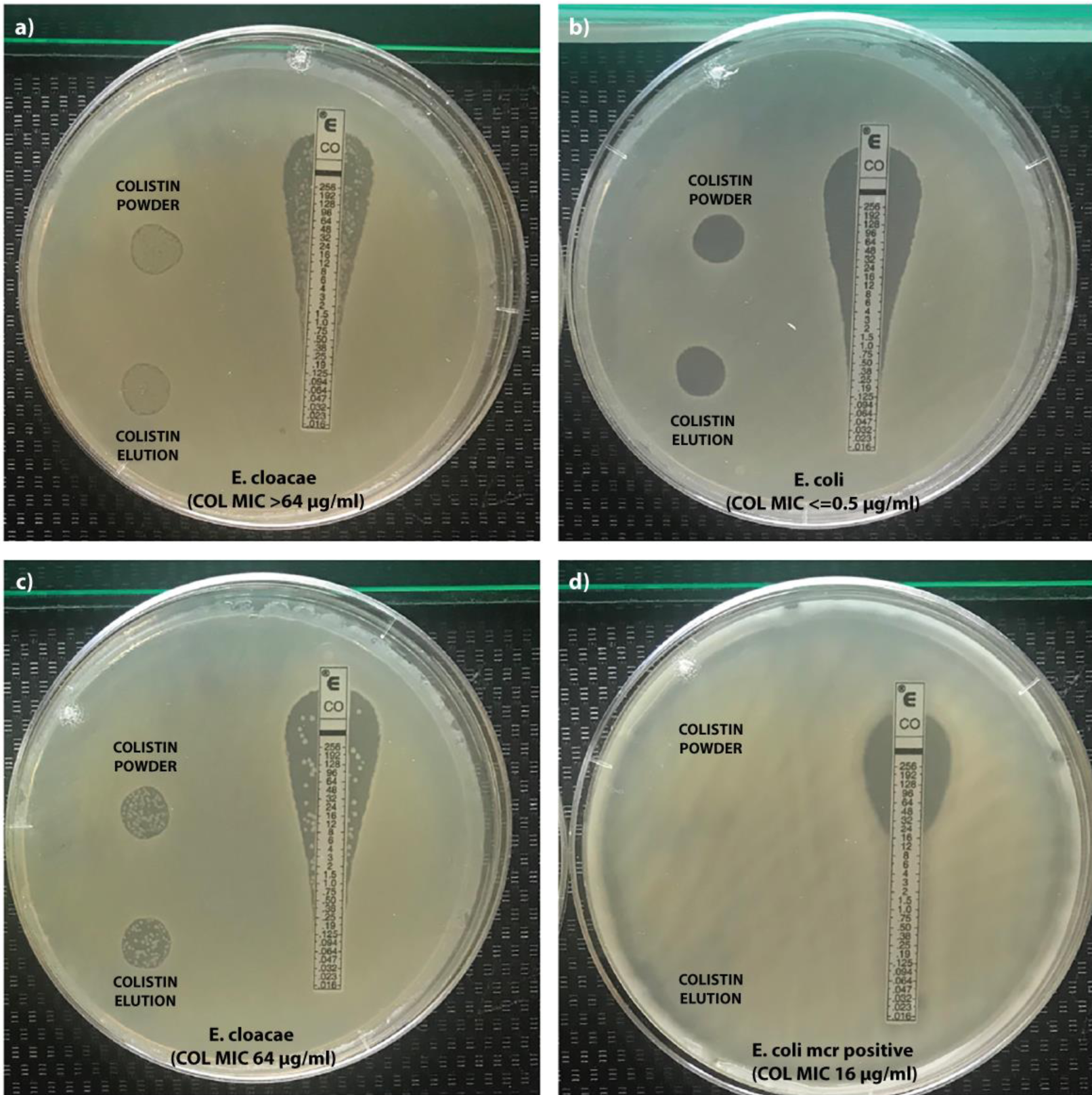
COL-DT. Examples of a colistin-resistant (a, c, d) and colistin-susceptible (b) strains by the drop test with 16 mg/L concentration of colistin in the 10 μL spot, obtained by powder solution or disk elution, as indicated. Strains a and c displyed well-defined colonies within the zone were categorized as resistant by the COL-DT. The species and the colistin broth microdilution MICs are indicated for each strain between brackets. The colistin gradient strip is shown only as indicative of the plane scale.

With a colistin solution made by elution of commercial disks in CA-MHB, we noted clear inhibition zones around the drop (a susceptible COL-DT) in 124/130 colistin-susceptible strains (drop zones range/median: 6-15/8.8 mm for *Acinetobacter* spp., 5-13/8.4 mm for *P. aeruginosa* and 7-11/9.6 mm for *Enterobacterales*). Again, 4 *E. coli mcr* producers susceptible by BMD grew without inhibition zone around the drop. In addition, 2 *P. aeruginosa* with MICs 2.0 μg/ml showed repeatedly zones with defined colonies or haze within the halo and were classified as resistant by COL-DT. All 141 colistin-resistant bacteria showed a colistin-resistant drop test result: 135/140 without an inhibition zone around the colistin drop and the remaining 5/140 (4 *E. clocacae* and 1 *A. baumannii*) showed repeatedly well-defined colonies invading the drop zone (Table 1S). With a polymyxin B solution made by elution of commercial disks in CA-MHB, we noted clear inhibition zones around the drop (a susceptible POL-DT) in 126/130 colistin-susceptible strains (drop zones range/median: 7-14/9.2 mm for *Acinetobacter* spp., 7-13/9.3 mm for *P. aeruginosa* and 9-14/9.8 mm for *Enterobacterales*). Exception were again 4 *E. coli mcr* producers (BMD MICs 2.0 μg/ml) that grew without inhibition zone around the drop (a resistant POL-DT). About 135/141 colistin-resistant bacteria, showed a resistant polymyxin B drop test result: 120/141 had no zone around the drop and 15/141 (mostly *Enterobacterales* and *Acinetobacter* spp.) showed repeatedly well-defined colonies invading the drop zone. The 6 strains exhibiting a false-susceptible POL-DT were 2 *P. aeruginosa*, 1 *E. coli mcr* producer, 1 *E. cloacae*, 1 *Samonella* spp, and 1 *Acinetobacter* spp (Table 1S).

The stored solutions showed identical performance to those freshly prepared. Zones around the drop remained unchanged over 12 months. Likewise, the repeatability studies showed agreement values (+/- 1 mm) in > 95% of the repetitions, without changes in the categorization of the strains.

#### 2) COL-AS

All 141 colistin-resistant bacteria grew in the spot plate irrespective of the concentration of colistin used (2.0 μg/ml or 3.0 μg/ml) or the brand of MHA tested (Difco and Britania).

Growth of a few colonies or semi-confluence growth was observed in 7/141, 5/141 and 4/141 with the 2.0 μg/ml in-house test, commercial brand and 3.0 μg/ml in-house test, respectively (Table 1S). Most semi-confluent growth was linked to *Salmonella* sp. (Figure 3).

**Figure 3.**
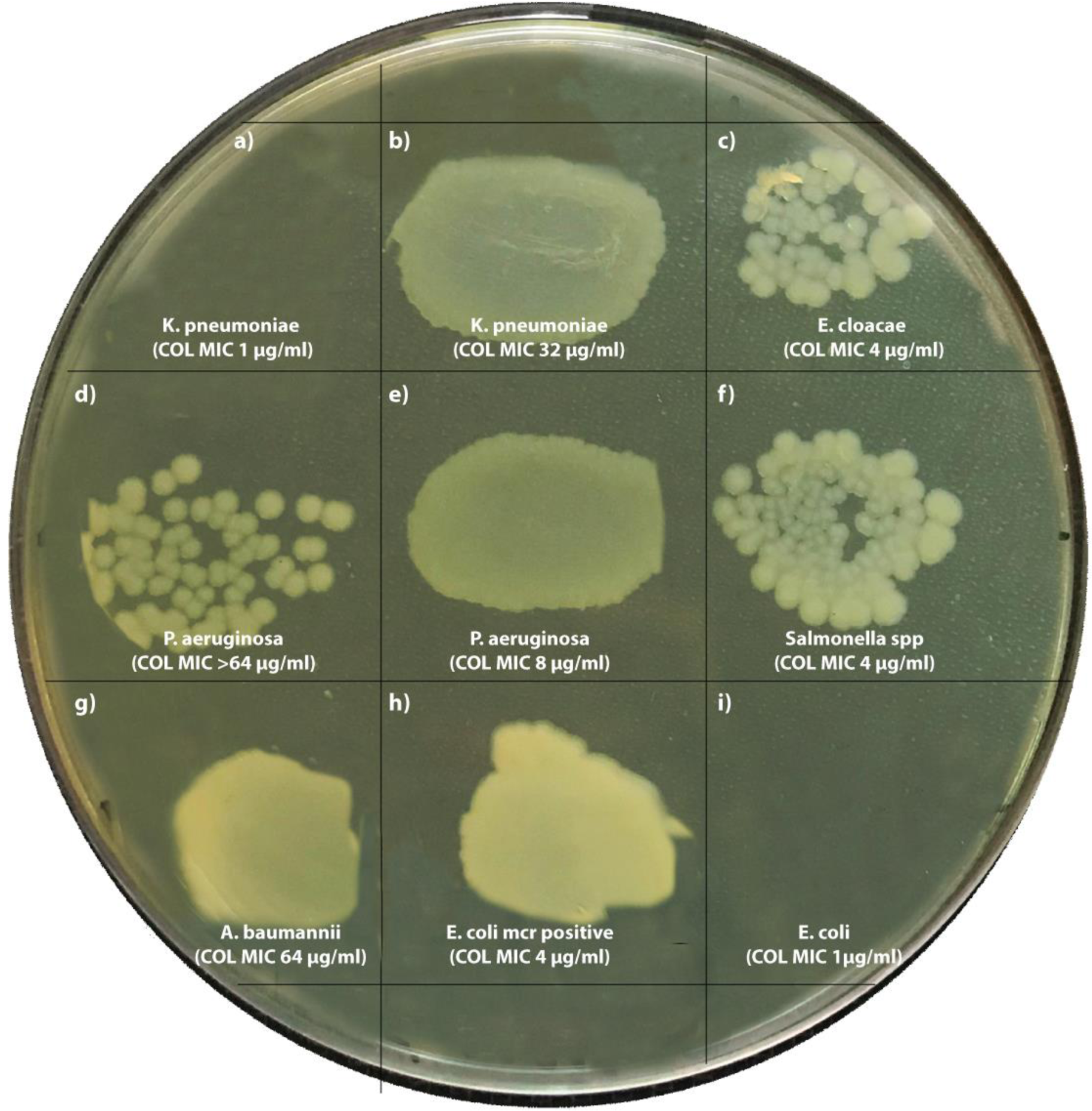
COL-AS. Examples of a colistin-resistant (b, c, d, e, f, g, h) and colistin-susceptible (a, i) strains by the in-house agar test with 3mg/L concentration of colistin. Strains c, d and f displayed semi-confluent growth. The species and the colistin broth microdilution MICs are indicated for each strain.

About 15/130 colistin-susceptible bacteria showed confluent or semi-confluent growth on the 2.0 μg/ml colistin plate. This number decreased to 5/130 when the concentration was increased to 3.0 μg/ml. These five strains corresponded to 3/4 *E. coli mcr* producers with a susceptible BMD MIC and 2 *Acinetobacter* spp. In the commercial method, 6 susceptible colistin strains grew on the COL-AS, the same 5 strains mentioned above along the remaining *E. coli mcr* producer with a susceptible BMD MIC (Table 1S).

The test repetitions performed in consecutive days, as well those performed with the stored plates up to 2 months, showed identical performance to those freshly prepared, without changes in the categorization of the evaluated strains nor affecting the type and amount of growth observed during storage.

#### Overall performance

The overall performance characteristics of the different colistin screening methods compared to reference BMD MIC are presented in Table 1 Considering that only the reference method (BMD), did not categorize all *mcr* producers as colistin resistant, but the other MIC methods did, performance was recalculated considering those *mcr* producers with discrepancies between MICs values using an consensual colistin MIC of 4 μg/ml, average of the values obtained by all three MIC methods.

**Table 1:** Error rates and categorical agreement produced by colistin screening tests for Gram-negative bacteria compared to reference broth microdilution or all MIC methods (recalculated values). The acceptable number of VME is a function of the number of resistant strains included in the validation, as recommended by the FDA.

| Test | Percentage of isolates with: |  |  | Criteria required for acceptance <sup>a</sup> |
| --- | --- | --- | --- | --- |
| Colistin screening test | categorical agreement (CA) and re-calculated CA (rCA) | very mayor errors (VME) and re-calculated VME (rVME) | mayor errors (ME) and re-calculated ME (rME) |  |
| COL-DT (colistin powder) | CA: 96.2<br>rCA:99.3 | VME: 0.7<br>rVME: 0.7 | ME: 3.1<br>rME: 0 | CA: >89.9<br>VME: <2.86<br>ME: <3.0 |
| COL-DT (colistin elution) | CA: 96,2<br>rCA:98.4 | VME: 0<br>rVME: 0 | ME: 3.8<br>rME: 1.6 |  |
| COL-DT (polymyxin elution) | CA: 94.2<br>rCA:96.6 | VME: 3.5<br>rVME: 3.4 | ME: 2.3<br>rME: 0 |  |
| COL-AS (2.0 µg/ml) | CA: 88.5<br>rCA:91.3 | VME: 0<br>rVME: 0 | ME: 11.5<br>rME: 8.7 |  |
| COL-AS (3.0 µg/ml) | CA: 95.5<br>rCA:97.7 | VME: 0.7<br>rVME: 0.7 | ME: 3.8<br>rME: 1.6 |  |
| ColTest (commercial agar spot) | CA: 95.4<br>rCA: 98.4 | VME: 0<br>rVME: 0 | ME: 4.6<br>rME: 1.6 |  |

a: according to FDA (Ref. 24).

## DISCUSSION

Colistin is the last resort agent used in the treatment of serious infections caused by multi-resistant *Enterobacterales, Pseudomonas aeruginosa* or *Acinetobacter* spp (1). Therefore, it is essential for laboratories to report accurate results with good essential agreement, being more important than for many other antimicrobial agents when it comes to patients with life-threaten infections. New therapeutic options such as ceftazidime-avibactam and meropenem-vaborbactam for extreme resistance bacteria, such as carbapenemase producers, have recently been approved and may be a reasonable alternative to colistin in the treatment of these life-threaten infections (25, 26). Despite having these antibiotics improved outcomes and a more predictable pharmacokinetic profile than colistin (27), low adoption has been reported in the US market with continued use of polymyxins (28). Until replacement of polymyxins by new, safer and more effective therapeutic options occurs, colistin susceptibility testing should be urgently improved.

Choosing an AST method for polymyxin susceptibility is challenging owing to poor penetration of drug in the agar medium, polycationic drug molecule binding to plates and changing MIC breakpoints by CLSI/EUCAST, as has happened in the last edition of CLSI (7), where only intermediate and resistant categories become available. A critical step in the validation processes of new or improved AST techniques, is the inclusion of representative strains, both susceptible and resistant, to avoid bias towards a positive or negative result. For this reason, the panel of strains included in this work was selected to guarantee half (whenever possible) of representative isolates with resistance to the agent to be tested, as suggested (22, 24). About 33%, 47% and 57% of the *P. aeruginosa, Acinetobacter* spp. and *Enterobacterales* of the panel, respectively, had colistin MICs > 2.0 μg/ml. It is also critical in the validation process to use the same bacterial inoculum across methods, since instability of the colistin resistant trait has been documented during the conservation, sub-culturing or thawing process of the bacterial frozen stocks (10, 29). For this reason, we evaluated each batch of strains using all methods on the same working day and the same bacterial suspension as inoculum.

In this work, we validated simple and low-cost methodologies, specially conceived for low- or middle-income countries, with limited access to reference BMD. Initially these methods do not replace MIC techniques, they would serve as screening to rule-out colistin-resistant strains from future analyzes. For this reason, methods included in this work have been designed to minimize the rate of VME, unlike the methods currently in use where this type of error is alarmingly high (9-14). If, at the same time, the proposed methodology presents a suitable ME rate, the screening tests could also contribute to rule-out those patients in whom a MIC-based colistin dose adjustment is not required.

The first methodology we recommend is the COL-DT, in which a single drop of colistin solution is applied on an inoculated agar surface. This technique was first used by Halling *et al*. who dropped aqueous solutions of defensins onto *Brucella* swabbed cultures plates (30). Subsequently, this methodology was adapted by Eric Jouy *et al*. to assess colistin susceptibility among *E. coli* of animal origin, including those with the *mcr*-1 gene (21). The authors using a colistin concentration of 8 mg/L in the 10 μL drop, were able to perfectly differentiate populations of *E. coli* with MIC <= 2.0 μg/ml from those with MIC > 2.0 μg/ml (21). When we extrapolated this methodology in species different from *E. coli*, especially *Klebsiella* and non-fermenters isolates from human sources, we observed haze or colonies within the drop zones in isolates with colistin MIC <= 2.0 μg/ml that hindered test interpretation. For this reason, a concentration of 16 mg/L in the 10 μL drop was further selected. Under these conditions, the COL-DT categorized as resistant all strains with MIC > 2.0 μg/ml. With respect to the strains with MIC <= 2.0 μg/ml, the drop test correctly categorized the panel of susceptible strains, with very few exceptions, as 4 *E. coli* strains *mcr* producers that were classified as resistant. These strains had a borderline MIC results, being susceptible by BMD but resistant by other reference methodologies, such as agar dilution or macro-dilution. The COL-DT, unlike the BMD, classified all the *mcr*-producing isolates as colistin-resistant. Remarkably, *E. coli* with BMD MIC of 2.0 μg/ml corresponded exclusively to isolates confirmed as producing *mcr*-1. Therefore, a colistin MIC value of 2.0 μg/ml obtained by BMD should be a call for additional testing, either phenotypic, such as COL-DT or COL-AS, or genotypic (PCR) to confirm the presence of *mcr* genes, especially among *E. coli*, as has been suggested by other authors as well (22). The COL-DT performance was re-calculated considering the average MIC value obtained by the 3 reference methods, and under these re-defined reference conditions, the drop test showed an improved ME rate.

Those laboratories that do not have access to colistin powder, will be able to obtain the solution from a commercial colistin disk by elution. Colistin was superior to polymyxin in performance, so it is recommended to use a colistin-based drop test. Use of commercial filter-paper disk as the source of the antibiotic (broth elution MIC) to overcome the cost and the limited access to powders, has been an old and widespread practice for anaerobic bacteria (31), and is now recommended for colistin MIC determination by CLSI (7). In this work, the disk eluted colistin solution had a performance equivalent to that of the powder. Subsequently, we evaluated the performance of the COL-DT using the BD Phoenix™ AST broth (Cat. 246003), ready-to-use MHB, cation-adjusted broth, with identical results as those presented in this work (not shown). The COL-DT offers additional advantages, even to other elution methods, namely: i) the solution is stable for more than 1 year in the refrigerator. Precautions must be taken to avoid contamination (periodical sterility controls, preferably monthly, by directly placing an aliquot in incubation at 35C, is required. ii) The COL-DT uses the same supplies as the diffusion method, widely used throughout the world (only a pipette or a 10-μl loop is required as additional material). Moreover, the drop test can be carried out on the same plate on which disks have been placed to carry-out the antibiogram, saving consumables. In this case, we recommend that the colistin drop be separated by at least 2 cm from the other placed disks; iii) Although it was not standardized for this purpose, the drop test has the potential to alert of possible sub-populations, a phenomenon frequently associated with *Enterobacter* and *Acinetobacter* spp. (32).

Numerous studies have demonstrated very good agreement between colistin agar dilution and BMD (34-38), with the exception of *P. aeruginosa* isolated from cystic fibrosis patients, in whom agar dilution might have more readily detected colistin resistance (34, 37). More recently, it was found that agar dilution was superior in terms of reproducibility and robustness, compared to broth dilution methods, for colistin MIC determination (38). Thus, in the second part of the study, we validate a single colistin concentration MHA plate for screening for colistin resistance. The technique is a slight variation of the method described in M07 (20), in which the antimicrobial dilution tested was limited to the concentration/s that encompass the interpretive breakpoints (between 2 – 4 μg/ml). We finally selected a concentration of 3.0 ug/ml which allowed to separate subpopulation with MIC <=2.0 μg/ml from those with MIC >2.0 μg/ml. Additional difference with the CLSI agar dilution method included the inoculum type and inoculation procedure: we used direct inoculation of the 0.5 Mc Farland turbidity suspension, swabbing a defined area of the plate (avg. inoculum of 1.0 × 10^6^ UFC/spot using a 10 μl loop for inoculation or 2.0 × 10^6^ UFC/spot by swabbing an 2 cm^2^ area). In the pre-standardization process, we evaluated inoculating the spot agar with a 1/10 dilution of the bacterial suspension adjusted to 0.5 Mc Farland, as recommended for dilution agar (20). This dilution caused false-susceptible results, specifically those strains that showed semi-confluent growth with the direct inoculation (not shown). Once again, the COL-AS was more robust in detecting *mcr*-producing strains than BMD. The commercial method (ColTest^®^) showed a performance equivalent to that developed in the laboratory, which could facilitate the adoption of this agar technique, especially in those facilities with limited access to colistin sulfate powder. The COL-AS had acceptable VME error (if any) and those ME were mainly due to those *E. coli mcr* producer that were categorized as susceptible by the BMD. The COL-AS also has the advantage for simultaneous evaluation of up to 12 or more isolates, even more if precaution of avoiding cross-contamination is taken. For this reason, a precautionary colistin resistant definition of > 1 colony was adopted.

Recently, an adaptation of the agar dilution method has been also recommended by CLSI as a test for colistin resistance for *Enterobacterales* and *P. aeruginosa*: this method uses MHA plates supplemented with serial dilutions of colistin (1.0 μg/ml, 2.0 μg/ml and 4.0 μg/ml) that are subsequently inoculated with 10 μl of a 1/10 dilution of a suspension of the tested bacteria adjusted to the turbidity of 0.5 of Mc Farland. The CLSI screening test showed no ME and only 0.5% of VME, exclusively among *Enterobacterales*, values comparable to those obtained with the COL-AS proposed here. An adaptation of the COL-AS proposed here was recently used for the specific detection of *mcr* producers, by adding an extra plate of colistin supplemented with EDTA, a well-known *mcr* inhibitor (39). Other agar-based colistin methods on the market were successfully utilized for screening purposes from clinical samples (40), showing the versatility of techniques based on agar dilution.

The limitations of this study were: 1) a limited access to non-fermenters colistin-resistant strains. These observations may justify more extensive validation of these methods on this group of pathogens; 2) methods were evaluated with a limited number of brands of disks and/or culture media, therefore, for users of other brands should be considered provisional until additional data demonstrating adequacy is available.

In conclusion, we have developed and validated methods (COL-AS, ColTest and COL-DT), suited to the systematic screening of resistance to colistin in gram negative bacilli with a performance similar to the reference BMD. These screening tests can be routinely performed in addition to the tests currently in use.

## ACKNOWLEDGEMENTS

We are indebted to those Institutions and professionals of the antimicrobial surveillance systems who referred the clinical isolates to the National Reference Laboratory for the repository collection of the NRL. We also acknowledge to Britania Argentina for gently providing ColTest for this study.

## TRANSPARENCY DECLARATIONS

Nothing to declare

